# NLGN3-tagging system for Evaluating Membrane Localization Applied to Drug Screening

**DOI:** 10.64898/2026.10.20.760290

**Authors:** Naoomi Tominaga, Kana Tominaga, Risako Okada, Takeshi Yamamoto

## Abstract

Aberrant membrane localization of proteins often results in functional impairment, contributing to the development of pathological conditions. Neuroligin**-**3 (NLGN3) is a rare but recurrent genetic risk factor for autism spectrum disorder (ASD). Comparing surface localization across NLGN3 variants requires that surface signal be referenced to expression level in the same cell. Using NLGN3 as a model, we developed I-CLiQ, a live-cell method in which the target coding sequence is introduced at a single site of a fixed vector backbone and DsRed is co-encoded as an internal expression reference, based on a 2A peptide of Perina nuda virus modified to abolish ribosomal skipping. Combining I-CLiQ with AlphaFold/AlphaMissense analysis, we evaluated ASD-associated NLGN3 variants. Structural modeling predicted steric clashes for p.R451C, p.P514S, and p.R597W. I-CLiQ-based flow cytometry confirmed that these variants significantly decrease surface expression. Furthermore, as a proof-of-concept application, we used I-CLiQ to evaluate two compounds, 4-phenylbutyrate (4**-**PBA) and 3-amino-1-propanesulfonic acid (3**-**APS); 4**-**PBA partially restored the surface expression of p.R451C and p.R597W. These findings establish I-CLiQ as a quantitative tool for analyzing membrane trafficking defects and demonstrate its applicability to compound screening for variants associated with conformational diseases.

## Introduction

Neuroligin**-**3 (NLGN3) is a postsynaptic cell adhesion molecule localized at both excitatory or inhibitory synapses ^1^, and plays a critical role in synapse formation and function ^2,3^. Genetic studies have identified multiple missense mutations in *NLGN3* in patients with autism spectrum disorder (ASD) ^4^. While some variants may directly perturb synaptic signaling, a substantial subset is predicted, on the basis of in silico and experimental analyses, to cause protein misfolding, leading to ER retention and reduced delivery to the cell surface—a mechanism commonly referred to as a trafficking defect ^5^. Elucidating how specific structural abnormalities influence trafficking efficiency is thus essential for understanding the pathophysiology of ASD and for informing the development of targeted therapeutic strategies ^6^.

Membrane proteins play pivotal roles in diverse cellular processes, including signal transduction, cell–cell adhesion, and ion transport ^7^. Consequently, mutations in genes encoding membrane proteins are frequently implicated in human diseases ^8^. To quantitatively evaluate the membrane localization of surface proteins, we previously developed an assay namely the 2A-assay that exploits the specific reactivity of the T2A peptide sequence derived from viruses ^9^. In this approach, the T2A sequence is fused to the C-terminus of the target protein, enabling sensitive detection of membrane-localized proteins using specific antibodies. The strategy of C-terminal tagging has demonstrated significant efficacy for membrane proteins that present their C-terminus to the extracellular environment. However, when this method is applied to type-I or -III membrane proteins, such as NLGN3, which possess an extracellular N-terminus and an intracellular C-terminus ^10^, an important limitation becomes evident. Numerous membrane proteins exhibit their N-terminus extracellularly, thus restricting the applicability of the 2A-assay to type-II proteins ^11^. The N-terminal domain constitutes the major functional region exposed to the extracellular milieu ^12,13^. Thus, accurate evaluation of N-terminal folding and surface presentation is essential for assessing their functional integrity ^14,3,15^. However, targeting the N-terminus for epitope tagging presents a significant technical challenge because of the presence of an N-terminal signal peptide (SP) ^16^. The SP is essential for cotranslational translocation into the endoplasmic reticulum (ER) but is typically cleaved by signal peptidase during or immediately after its translocation ^17,18^. Consequently, simple fusion of an epitope tag to the extreme N-terminus often results in removal of the tag together with the SP or, alternatively, interferes with SP recognition and thus disrupts proper membrane insertion ^17^. Because of these difficulties, a broadly applicable N-terminal labeling strategy that enables quantitative assessment of membrane localization in cells would be highly valuable ^19^.

In this study, we established an N-terminal tagging strategy intended for quantitative assessment of the membrane localization of type-I transmembrane proteins such as NLGN3 as a proof of concept. We term this approach the “I-CLiQ” (type-I cell membrane protein localization quantitative) method. By combining this experimental platform with in silico structural analyses using AlphaFold ^20^ and pathogenicity prediction using AlphaMissense ^21,22^, we systematically predicted the structural and functional impacts of NLGN3 variants. Furthermore, as a proof of concept, we applied the I-CLiQ system to assess the efficacy of 4-phenylbutyrate (4-PBA) and 3-amino-1-propanesulfonic acid (3-APS)^23^ as chemical chaperones, demonstrating its potential utility for evaluating compounds that modulate plasma membrane trafficking of type-I membrane proteins. These findings support the utility of I-CLiQ for characterizing variant-associated alterations in cell membrane localization and assessing their responses to pharmacological intervention.

## Results

### Structural validation of the NLGN3 isoform and rationale for pathogenicity analysis

To establish a structural basis for our study, we first predicted the three-dimensional model of the specific NLGN3 isoform used in our experiments (Q9NZ94-2) using ColabFold (AlphaFold2), as no experimental structure is available for this isoform. The prediction was performed for the monomer. We then compared this model with the AlphaFold Database model AF-Q9NZ94-F1-v6. First, amino acid sequence alignment confirmed that the two isoforms share 98% sequence identity (Table S1), ensuring consistent residue numbering. Second, we compared the two predicted models. The sequence coverage of the multiple sequence alignment (MSA) is shown in Figure S1A. The predicted lDDT values along the sequence are plotted in Figure S1B, which illustrates the residue-wise relationship between the prediction confidence and sequence position. The predicted aligned error (PAE) for Q9NZ94-2 was also calculated (Figure 1A, and S1C), and its pattern closely resembled the PAE of AF-Q9NZ94-F1-v6. The analysis indicated that the N- and C-terminal regions were predicted to be disordered, with low pLDDT scores (Figure 1B). Additionally, superimposition of the predicted Q9NZ94-2 model onto AF-Q9NZ94-F1-v6 gave a root mean square deviation (RMSD) of 0.857 Å (Figure 1C, and S1D).

**Figure 1.**
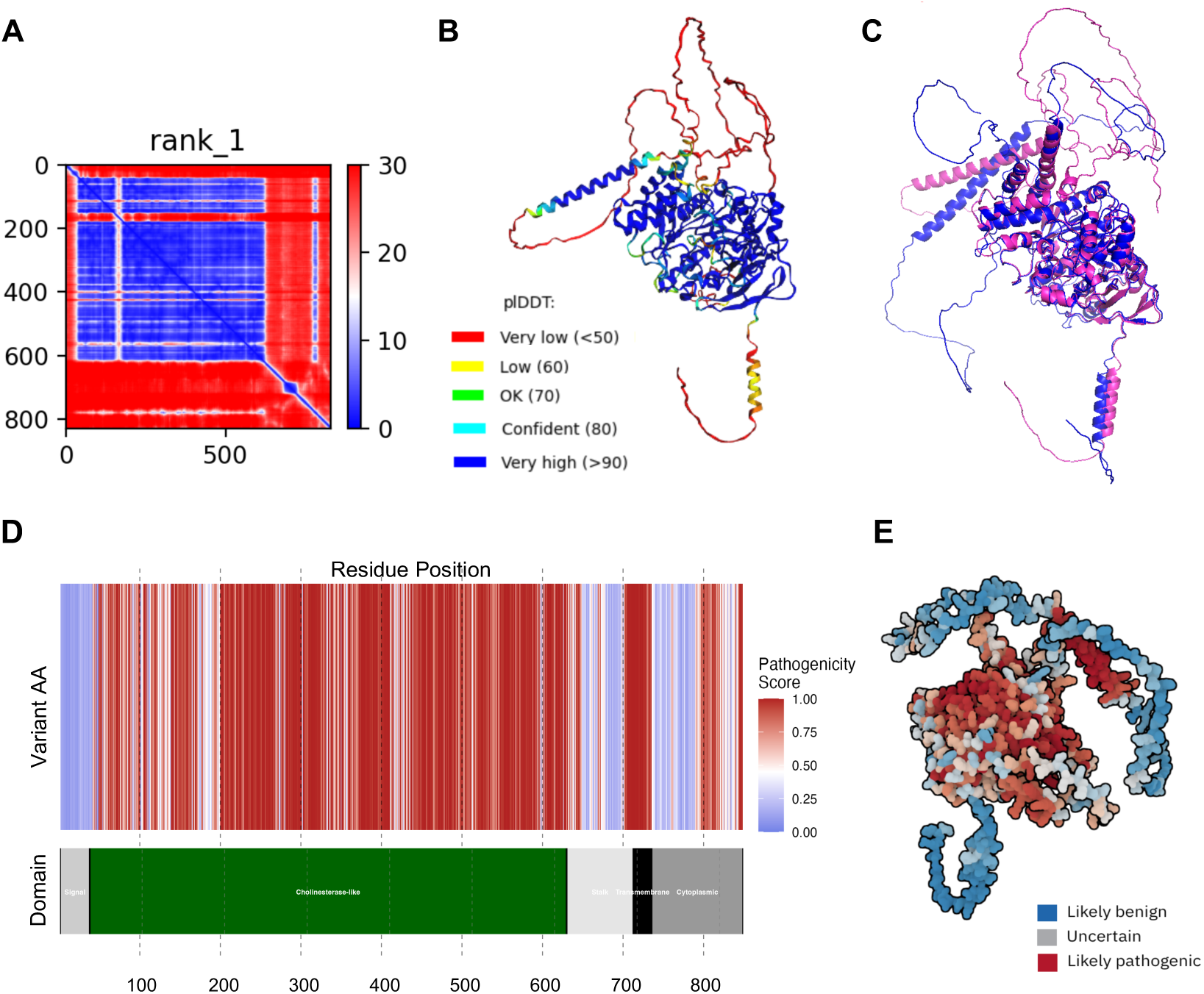
Structural modeling and pathogenicity prediction of NLGN3 using AlphaFold and AlphaMissense. All predictions were performed for the NLGN3 monomer. **(A)** Predicted Aligned Error (PAE) plot for the NLGN3 Q9NZ94-2 model generated by AlphaFold, indicating the expected error in the relative positioning of domains. **(B)** Predicted model colored by per-residue pLDDT scores (blue: high confidence; yellow: low confidence). **(C)** Superimposition of the AlphaFold-predicted model of Q9NZ94-2 (rank 1) (color) and the AlphaFold Database model AF-Q9NZ94-F1-v6 (gray). The RMSD between the two models is 0.857 Å. **(D)** Landscape of pathogenicity predictions by AlphaMissense. (Upper) Distribution of variants predicted to be pathogenic (red) or benign (blue). (Lower) Domain organization of NLGN3. **(E)** Mapping of AlphaMissense pathogenicity scores onto the predicted model. Variants predicted to be pathogenic are clustered within the folded core domains.

The high sequence identity and the close correspondence between the two predicted models support the use of the AlphaMissense dataset, which is based on the canonical isoform, for our target isoform. Utilizing the AlphaMissense data, we mapped the pathogenicity scores onto the model (Figure 1D). The information in Figure 1D is further visualized as a three-dimensional representation in Figure 1E. Variants predicted to be pathogenic (red) are densely clustered within the folded core domains (Figure 1E). Together, these analyses suggested that the folded core domains and the transmembrane domain are particularly vulnerable to missense mutations. Since mutations disrupting the structural core often trigger quality control mechanisms leading to endoplasmic reticulum (ER) retention ^5,8,14^, these findings underscore the necessity of developing a method to quantitatively evaluate the membrane trafficking efficiency of these pathogenic variants.

### In silico pathogenicity profiling and structural impact analysis of selected variants

To further investigate the molecular basis of NLGN3 dysfunction, we selected five representative disease-associated missense variants previously reported in clinical studies (Table 1). First, we mapped these variants onto the AlphaMissense landscape (Figure 2A). The p.R451C, p.P514S, and p.R597W variants exhibited high pathogenicity scores (over 0.564) and were located within the highly conserved cholinesterase-like domain, supporting their functional importance. On the other hand, p.V341A (0.514) falls into the uncertain-significance range, and p.T632A (0.06) is predicted to be more likely benign. With the exception of p.T632A, the variants were classified as deleterious or (possible) damaging by SIFT and PolyPhen**-**2 (SIFT scores ≈ 0, PolyPhen**-**2 scores 0.7–1.0). In contrast, p.T632A was predicted to be tolerated and benign by both methods (SIFT 0.08, PolyPhen-2 0.007), consistent with its low AlphaMissense score.

**Table 1.** Five NLGN3 mutants predicted to be pathogenic.

| Variant ID | protein change | WT | Mutant | AlphaMissense | SIFT | Polyphen2 |
| --- | --- | --- | --- | --- | --- | --- |
| rs749067360 | p.V341A | gta | gCa | 0.5143 | 0.00 | 0.733 |
| rs121917893 | p.R451C | cgt | Tgt | 0.9417 | 0.00 | 1.000 |
| rs1569485503 | p.P514S | cct | Tct | 0.9865 | 0.00 | 0.986 |
| rs878853147 | p.R597W | cgg | Tgg | 0.9942 | 0.00 | 1.000 |
| rs144914894 | p.T632A | acc | Gcc | 0.0605 | 0.08 | 0.007 |

**Figure 2.**
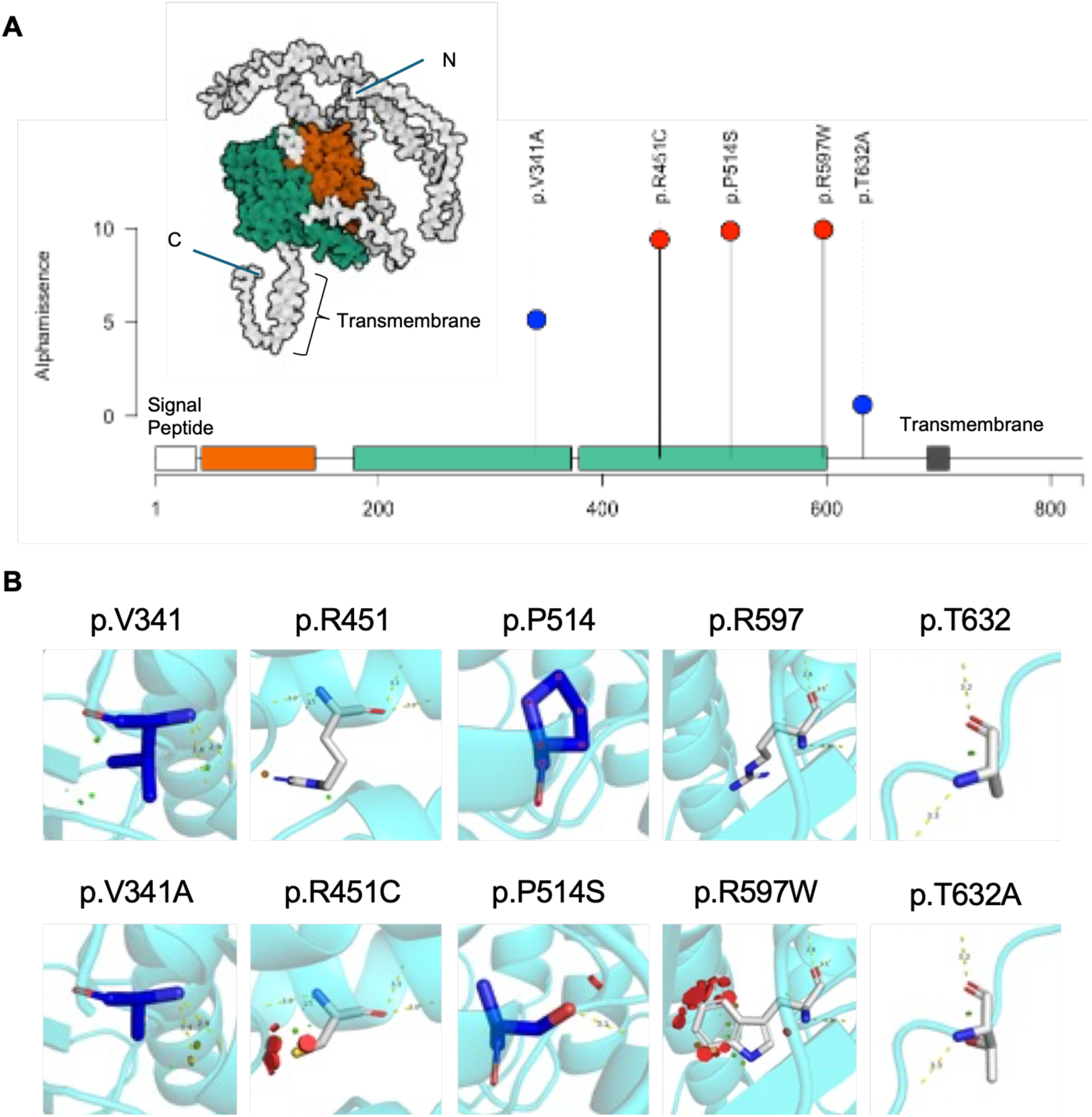
Structural mapping and consensus pathogenicity prediction of disease-associated NLGN3 variants. **(A)** Lolliplot visualizing the landscape of AlphaMissense pathogenicity scores for NLGN3. The x-axis represents the amino acid position, and the y-axis indicates the predicted pathogenicity score. The positions of the five selected variants are marked. The predicted 3D model shows the spatial arrangement of the functional domains and their relative positions along the polypeptide chain. **(B)** Structural simulation of the specific variants using the PyMOL Mutagenesis Wizard. The upper panels display the WT local environment, while the lower panels show the corresponding variant models. In the variant structures (bottom), red discs indicate van der Waals overlaps between the introduced side chain and neighbouring atoms in the modelled rotamer.

Next, to assess the structural impact of these substitutions, we performed in silico mutagenesis analysis using PyMOL (Figure 2B). Comparison of the mutant models (lower panels) with their wild-type (WT) counterparts (upper panels) revealed distinct structural consequences. The substitutions p.R451C, p.P514S, and p.R597W resulted in the appearance of prominent red discs in the mutant models, which indicate van der Waals overlap between the mutated side chain and the surrounding atoms in the modelled rotamer. These findings suggested that these specific rotamers may be energetically and physically unfavorable within the native backbone conformation, which could destabilize the protein structure or necessitate considerable local remodeling. In contrast, the other two variants displayed minimal steric conflicts, suggesting that they may compromise NLGN3 function through mechanisms distinct from direct steric strain. These two approaches showed a high degree of concordance, with p.R451C, p.P514S and p.R597W classified as deleterious by both AlphaMissense and in silico mutagenesis analysis. However, p.V341A, which resides within the cholinesterase-like core domain, may have clinical relevance that is not fully captured by structure-based predictions alone. In addition, p.T632A, despite its benign-leaning AlphaMissense score, could still exert context-dependent effects and thus warrants cautious interpretation in functional and clinical studies.

### Establishment of the I-CLiQ System for Quantification of Plasma Membrane Localization

To experimentally validate the membrane localization of these variants, we developed the I-CLiQ system using a specialized epitope tag (Figure 3A). As an initial proof of concept, we first evaluated the p.P514S variant using this system. The I-CLiQ tag is derived from the 2A peptide sequence of Perina nuda virus. Unlike the parental 2A peptide, which induces ribosomal skipping, we introduced four point substitutions (D13H, V16I, G21V and P22F; see Methods) to abolish this activity, allowing the sequence to function as a stable epitope tag that is genetically fused to the N-terminus of NLGN3. The substitutions at positions 21 and 22 disrupt the C-terminal NPGP motif required for ribosomal skipping, so that skipping is abolished while the antigenic region of the peptide is retained.

**Figure 3.**
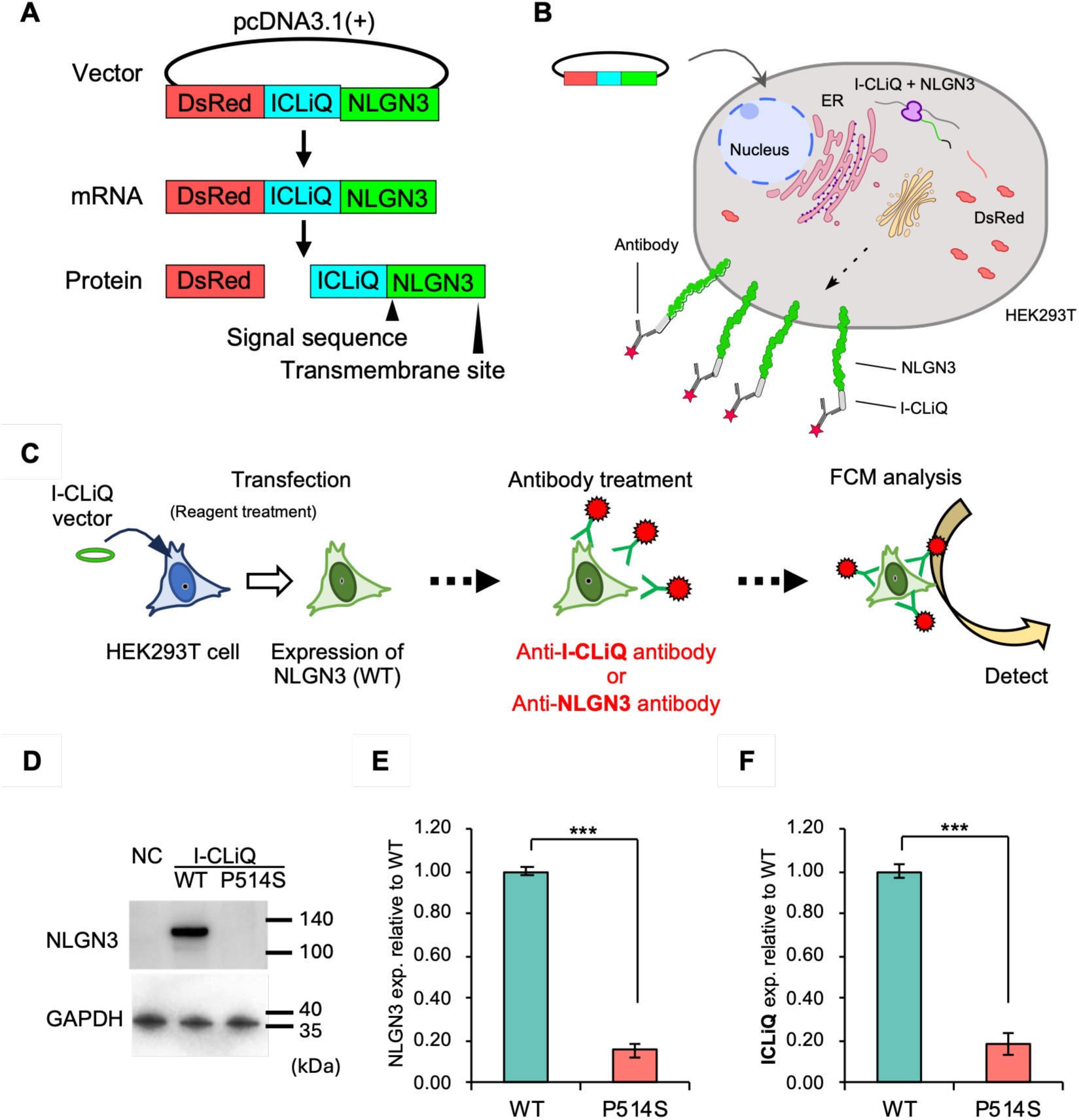
Establishment and validation of the I-CLiQ-tagged NLGN3 expression system. **(A)** Schematic representation of the gene expression cassette, comprising DsRed2, the I-CLiQ tag and human NLGN3 isoform 2 in a single open reading frame. The native signal peptide of NLGN3 lies at the N-terminus of the NLGN3 segment. **(B, C)** Schematic model of the surface display assay using flow cytometry (FCM). HEK293T cells expressing the construct are stained with an anti-I-CLiQ antibody to detect membrane-localized NLGN3 fusion proteins. **(D)** Western blot analysis of HEK293T cell lysates. Expression of the construct was examined using an anti-NLGN3 antibody. Anti-GAPDH was used as a loading control. N.C., untransfected cells. **(E, F)** Quantitative evaluation of cell surface expression by FCM. Cells were stained with either anti-NLGN3 antibody (E; n = 4 independent experiments) or an anti-I-CLiQ antibody (F; n = 3 independent experiments), each averaged from 3 technical replicates. Surface expression values were normalized to co-expressed intracellular DsRed2 and are expressed relative to WT. Data are shown as mean ± SEM. Statistical analysis was performed using an unpaired two-tailed Student’s t-test. ***; *p* < 0.001.

Next, to verify whether the I-CLiQ tag allows for the quantitative assessment of membrane trafficking, we established a robust flow cytometry protocol (Figure 3B, 3C). To ensure maximal sensitivity and specificity, we optimized the staining parameters using live cells without permeabilization, thus exclusively detecting cell-surface proteins. We determined that an antibody incubation time of 2 h (Figure S2A and S2B) and an anti-I-CLiQ antibody concentration of 37 μg/mL (1:25 dilution of the stock) provided the optimal signal-to-noise ratio (Figure S2C and S2D). Using these optimized conditions, we evaluated the surface expression of the construct. To enable quantitative comparison across constructs and experiments, surface I-CLiQ signal was normalized on a per-cell basis to co-expressed intracellular DsRed, which served as an internal reference for transfection efficiency and expression level. Because transfection efficiency varies and not all cells express the construct, the analysis was restricted to DsRed-positive live cells by gating, so that the quantification was performed exclusively on the expressing population. Normalized values are expressed relative to WT (set to 100%). This DsRed-based normalization allows relative surface expression to be compared across variants independently of differences in transfection or total expression.

We next examined expression by western blotting of whole-cell lysates with an antibody against NLGN3 (Figure 3D). A band at the expected molecular weight of approximately 94 kDa was detected for the WT construct. Because whole-cell lysates report the total protein pool rather than the surface pool, all quantitative comparisons of surface localization were made by live-cell flow cytometry of non-permeabilized cells. Relative to the WT, the surface signal of p.P514S was reduced to 15.4% when measured with the anti-NLGN3 antibody (Figure 3E) and to 18.5% when measured with the anti-I-CLiQ antibody (Figure 3F). Retention of the I-CLiQ epitope on surface-exposed molecules is supported by this agreement between the two antibodies, together with the detection of a specific anti-I-CLiQ signal on transfected, non-permeabilized cells and its absence on non-transfected cells (Figures S2A). These results demonstrate that the I-CLiQ tag is accessible on the cell surface and that the anti-I-CLiQ antibody serves as a robust tool for evaluating membrane localization.

The I-CLiQ workflow is based on transient transfection, with per-cell DsRed normalization and gating of the DsRed-positive population serving to accommodate variation in transfection efficiency. To assess whether this transient-transfection format could be extended to a neuronal cell line, we quantified DsRed-positive cells by flow cytometry (Figure S3). In HEK293T cells, approximately 10% of cells were DsRed-positive (Figure S3A). In SK-N-SH cells, the DsRed-positive fraction remained below 0.1% across the DNA-to-reagent ratios tested with PEI-MAX (Figure S3B) and reached only approximately 0.4% with Lipofectamine 3000 (Figure S3C). The expressing population was thus too small for reliable gating, indicating that applicability of the present transient-transfection format is constrained in this neuronal cell line.

### Pathogenic mutations impair membrane localization

Building upon the structural predictions and the validation of the I-CLiQ system, we experimentally investigated the degree of membrane localization of the five selected pathogenic variants. First, we quantified the cell surface abundance of each variant using the I-CLiQ flow cytometry assay (Figure 4A). Consistent with AlphaMissense and the in silico prediction of structural instability, the p.R451C, p.P514S, and p.R597W variants exhibited substantial decreases in surface expression. Compared with the WT, the variants resulted in decreases in surface expression to 30.3%, 25.8%, and 32.9%, respectively (*p* < 0.001). The p.V341A variant (AlphaMissense pathogenicity score: 0.514), which had an AlphaMissense pathogenicity score close to the high-risk threshold (0.564), exhibited a membrane localization level comparable to that of the three strongly pathogenic variants in the I-CLiQ assay, representing 38.3% of the WT. In contrast, the p.T632A variant exhibited an even higher level of membrane localization (134.3%) than the WT. We therefore used the value obtained for p.V341A, the highest surface expression among the variants reported as pathogenic in previous studies, as a dataset-specific reference threshold for reduced surface expression (38.3% of the WT). This value applies to the present dataset and is not proposed as a diagnostic cut-off. As p.T632A illustrates, pathogenicity cannot be defined by reduced surface localization alone.

**Figure 4.**
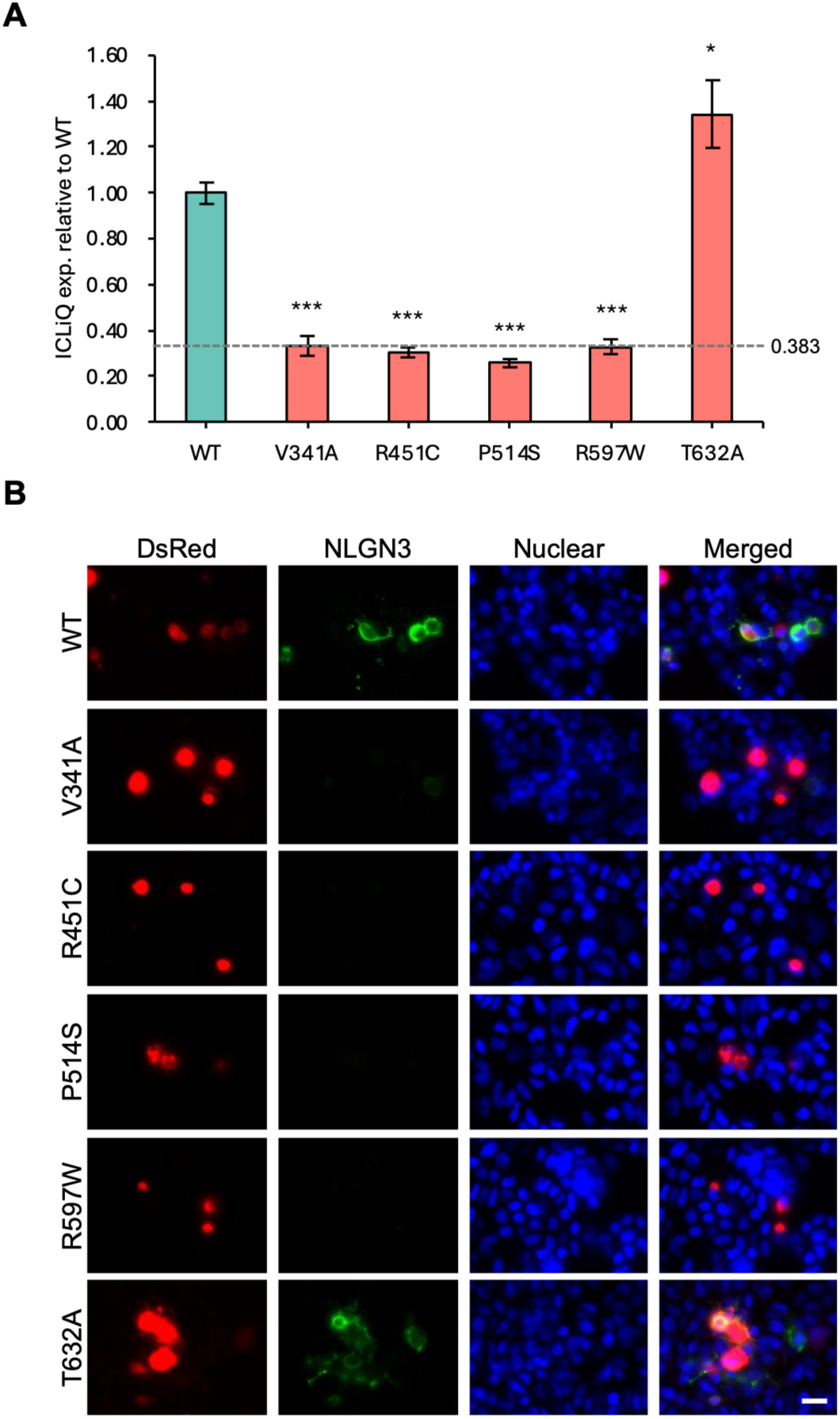
Quantitative and microscopic assessment of cell surface localization of NLGN3 variants. **(A)** Quantitative analysis of cell surface expression using the I-CLiQ system. Surface expression values were normalized to co-expressed intracellular DsRed2 and are expressed relative to WT. Data are mean ± SEM, n = 3 independent experiments. Statistical analysis was performed using one-way ANOVA followed by Tukey’s honestly significant difference test. \**P* < 0.05, \*\*\**p* < 0.001 (vs. WT). **(B)** Immunofluorescence of fixed, non-permeabilized HEK293T cells expressing NLGN3 variants, stained with an anti-NLGN3 antibody so that only surface-exposed NLGN3 is detected. Images were acquired with a 100x objective. Negative controls are shown in Figure S3. Scale bar, 20 μm.

To corroborate these findings visually, we performed immunofluorescence staining of fixed, non-permeabilized cells using an anti-NLGN3 antibody, which detects only surface-exposed NLGN3 (Figure 4B). The specificity of the staining was confirmed by negative controls (Figure S4). WT cells showed a continuous signal outlining the cell periphery, whereas p.V341A, p.R451C, p.P514S and p.R597W showed a markedly weaker peripheral signal. p.T632A retained a peripheral signal at least as strong as that of the WT. These patterns were consistent with the relative surface expression levels measured by the I-CLiQ assay. Collectively, these data demonstrate that the I-CLiQ assay reliably quantifies the membrane localization of NLGN3 variants. They also indicate that experimental assessment of membrane localization is essential, as the results can differ from those of pathogenicity predictions based solely on structural modeling.

### Evaluation of chemical chaperone efficacy using the I-CLiQ screening platform

Cells expressing the NLGN3 variants were treated with 1 mM 4-phenylbutyrate (4**-**PBA) or 10 μM 3-amino-1-propanesulfonic acid (3-APS) for 24 h, and their surface expression was quantified relative to that of DMSO-treated WT cells (Figure 5). This assay successfully detected basal surface levels, and 4**-**PBA treatment induced a significant increase in the membrane localization of the p.R451C and p.R597W variants. With respect to p.R451C, membrane localization increased from 0.306 ± 0.014 with DMSO treatment to 0.364 ± 0.033 with 4**-**PBA treatment, and with respect to p.R597W, it increased from 0.412 ± 0.019 to 0.537 ± 0.045. Compared with DMSO-treated cells, 4**-**PBA treatment induced a significant decrease in the membrane localization of the p.P514S and p.T632A variants. Moreover, in the 3-APS-treated groups, NLGN3 membrane localization was reduced for all the variants. This assay thus enabled the quantification of membrane localization and its application to the evaluation of compounds that modulate surface expression.

**Figure 5.**
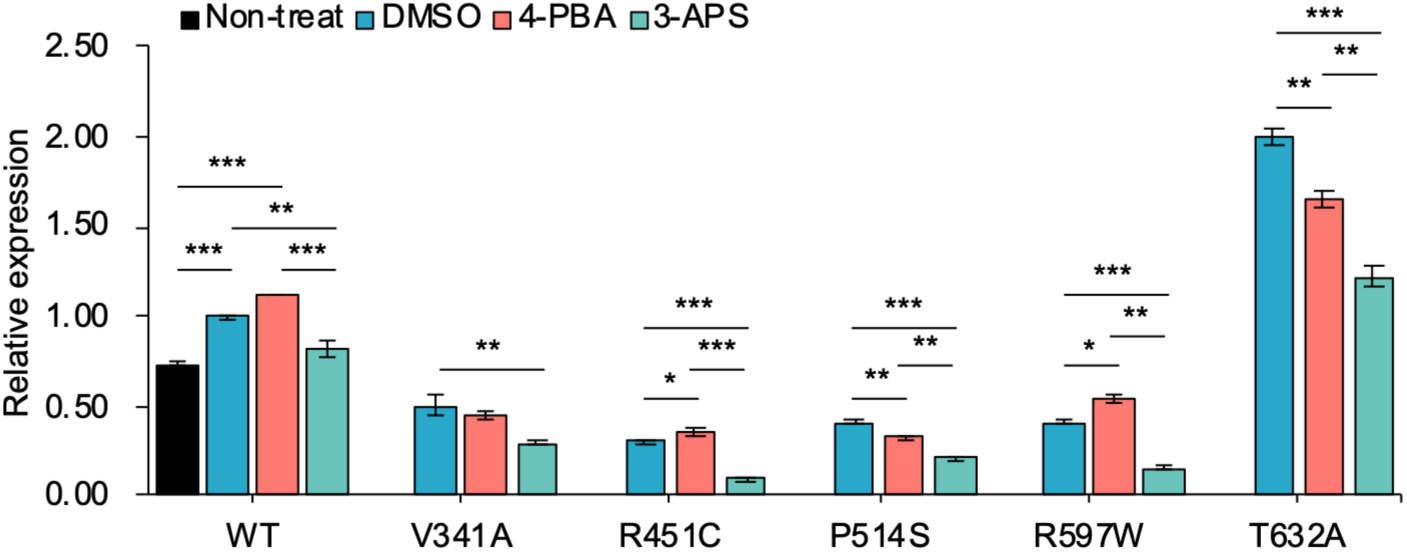
Assessment of the effect of 4-PBA or 3-APS treatment on the surface localization of NLGN3 variants. HEK293T cells expressing WT or variant constructs were treated with 4**-**PBA (1 mM), 3**-**APS (10 μM) or vehicle (DMSO) for 24 h. Surface signal was normalized to co-expressed intracellular DsRed2 and is expressed relative to the DMSO-treated WT control (set to 1.0). Data are means ± SEM. of n = 3 independent experiments. Statistical analysis was performed using one-way ANOVA followed by Tukey’s honestly significant difference test, comparing each treatment with the DMSO control within each construct. * *p* < 0.05, ** *p* < 0.01, *** *p* < 0.001.

## Discussion

In this study, we developed an N-terminal tagging system for type-I membrane proteins, in which the tag is placed upstream of the signal peptide, and used it to quantify membrane localization for NLGN3. This approach was employed to screen potential therapeutic compounds aimed at rectifying localization defects in ASD-associated NLGN3 variants. By introducing the “I-CLiQ” system, an N-terminal tagging strategy in which the tag is retained on a detectable fraction of the translated molecules, we quantified the pronounced reduction in membrane localization observed for NLGN3 variants carrying substitutions within the core domain. While C-terminal tagging methods, including our previously developed 2A-based assay, are effective for type-II membrane proteins ^9^, they are inherently limited for type-I membrane proteins because they cannot verify whether the extracellular N-terminal domain is correctly processed and presented on the cell surface. Conventional N-terminal tagging strategies are also poorly suited to the specific quantification of surface localization. Fluorescent protein tags, such as GFP, report the total fluorescence pool, including intracellularly retained, ER/Golgi-resident, and aggregated proteins; they are therefore informative for overall expression but cannot, by themselves, isolate the membrane-localized fraction. Short epitope tags, such as HA, when placed at the extreme N-terminus upstream of the signal peptide, are often liable to be removed together with the signal peptide during processing, which can lead to loss or heterogeneity of the surface signal. However, placing an epitope tag immediately downstream of the signal peptide may still affect signal peptide cleavage, as well as the folding, glycosylation, cell-surface trafficking, and function of the target protein. Moreover, an epitope tag alone is not well suited for normalizing intracellular expression levels or enabling reliable comparisons across samples. Detection relies on antibody binding to the tag under non-permeabilized, live-cell conditions, ensuring that internalized, retained, or aggregated proteins are excluded from the quantification. This provides a more specific and accurate measure of the membrane-localized fraction than total-signal or permeabilization-based readouts. The I-CLiQ system addresses these limitations by using a modified 2A-derived tag fused to the N-terminus, which is retained on a subset of the translated molecules and thereby provides a detectable epitope at the cell surface. This allows for rapid, live-cell quantification of functional surface expression with high sensitivity and specificity, as demonstrated by the comparable detection levels between anti-I-CLiQ and anti-NLGN3 antibodies. Cross-validation was possible here because a well-characterized antibody against the extracellular domain of NLGN3 is commercially available (Figures 3E and 3F); for many type-I membrane proteins such antibodies remain limited, and their epitopes can be sensitive to disease-associated mutations or conformational changes. By targeting an invariant N-terminal tag, the anti-I-CLiQ antibody provides a route to surface quantification that does not depend on the availability of a target-specific antibody. The I-CLiQ tag is positioned N-terminal to the signal peptide, and the anti-I-CLiQ antibody therefore reports the population of molecules on which the tag is retained. That this population provides a faithful readout of surface NLGN3 is supported by the concordance between the anti-I-CLiQ and anti-NLGN3 measurements across the variants in which both could be assessed (Figures 3E and 3F), since a variant-dependent difference in tag retention would be expected to disrupt this agreement.

By combining computational structural modeling with cellular validation, this study identified an association between mutation-induced structural destabilization predicted in silico and defects in protein trafficking measured experimentally. R451C ^24^, P514S ^25^, and R597W ^25^ have previously been shown to be retained in the ER, and our analyses using AlphaFold and AlphaMissense suggested that steric clashes (van der Waals overlaps) may contribute to these defects, although the structural basis was inferred from modeling and was not directly demonstrated. On the other hand, although V341A was predicted by computational structural modeling to have a low mutagenic effect, both clinical evidence and the results of the I-CLiQ assay indicated that this variant is indeed pathogenic. In addition, T632A was predicted to be relatively benign by structural modeling but exhibited increased expression in cell-based assays. Importantly, increased surface localization should not be equated with a benign functional outcome. In NLGN3, the autism-associated R451C mutation reduces protein levels yet enhances excitatory synaptic transmission through a gain-of-function mechanism, indicating that the relationship between NLGN3 abundance, surface localization, and synaptic function is not strictly proportional ^26^. In this context, the increased surface localization observed for p.T632A may not necessarily be functionally neutral and could reflect a distinct mode of dysregulation rather than a simple loss of pathogenicity. This interpretation remains speculative, however, and direct functional validation will be required to determine its physiological consequences. Collectively, these findings are consistent with a “conformational disease” model for these NLGN3 variants, in which the principal pathogenic mechanism is impaired trafficking rather than a primary defect in molecular function or ligand binding. Moreover, they highlight that although structural modeling provides critical mechanistic insight, certain pathogenic variants can be revealed only by cell-based functional assays, underscoring the importance of integrating both approaches to bridge this gap in variant interpretation.

The I-CLiQ system also demonstrated its potential utility for evaluating compounds that modulate NLGN3 surface expression. The two compounds evaluated in this study act through distinct mechanisms. 4-PBA is a chemical chaperone that assists protein folding and promotes the trafficking of misfolded proteins to the cell surface. In contrast, 3-APS (tramiprosate) was originally developed as an anti-aggregation agent that binds soluble amyloid-β and maintains it in a non-fibrillar state, thereby inhibiting its aggregation ^27,28^. Given that misfolded proteins are prone to aggregation, the established anti-aggregation activity of 3-APS suggested that it might also limit the aggregation of misfolded NLGN3 variants and thereby support their surface delivery. We therefore evaluated 3-APS alongside 4-PBA as a mechanistically distinct candidate. 4-PBA significantly increased NLGN3 membrane localization for the p.R451C and p.R597W variants. Several lines of evidence indicate that chemical chaperones can modulate the folding and trafficking of neuroligin family proteins, including autism-associated NLGN4X variants for which 4-phenylbutyrate (4-PBA) partially restores synaptic function ^29^. In addition, a lipophilic 4-PBA derivative prevents pathological protein aggregation ^23^, and ER stress in autism spectrum disorder has been linked to NLGN3 and related synaptic adhesion molecules, with 4-PBA proposed as a prototypical chemical chaperone to alleviate such proteostatic stress ^30^. 3-APS, in contrast, reduced surface localization across all variants tested. Because chemical chaperones and anti-aggregation compounds act on the specific conformational state of each mutant, a uniform, beneficial effect across structurally distinct variants would not necessarily be expected, and the divergent responses observed here likely reflect differences in the underlying conformational defects of individual variants. The molecular basis of these variant- and compound-specific responses remains to be fully elucidated and will require dedicated mechanistic studies. These findings nonetheless illustrate that the I-CLiQ assay can resolve differential, variant-specific responses to candidate compounds, supporting its utility for evaluating trafficking modulators. Whether such compound-induced changes in surface localization are clinically meaningful remains to be determined and will require further functional analysis.

In conclusion, we have established the I-CLiQ system as a practical tool for quantifying the surface localization of NLGN3, a type-I membrane protein. Among the five variants examined, reduced surface localization coincided with predicted local steric effects for p.R451C, p.P514S and p.R597W, but not for p.V341A, which showed reduced localization despite a low predicted steric effect, or for p.T632A, which showed increased localization despite a benign-leaning prediction. Structure-based prediction alone therefore did not account for the surface-localization phenotypes observed here, and cell-based measurement was required to resolve them. The I-CLiQ system provides such a measurement and, as shown here, a platform for evaluating compounds that modulate surface trafficking, with potential relevance to conformational diseases.

## Methods

### ColabFold

The three-dimensional structure of NLGN3 (Q9NZ94-2) was predicted using ColabFold (ColabFold v1.5.5: https://github.com/sokrypton/ColabFold), which provides an implementation of AlphaFold2 with accelerated MMseqs2-based multiple sequence alignment generation ^20^. For each sequence, ColabFold generated five structural models ranked by the predicted local distance difference test (pLDDT), and the top-ranked model (rank_001) was used for subsequent analyses, following the original ColabFold/AlphaFold recommendations. Predictions were performed for the NLGN3 monomer. NLGN3 is known to form dimers, and the effect of dimerization on the local environment of the residues analyzed here was therefore not assessed. In the predicted models, per-residue pLDDT scores (0–100) were used as a local confidence metric: pLDDT > 90 was interpreted as very high confidence, pLDDT between 70 and 90 as high confidence, pLDDT between 50 and 70 as low confidence, and pLDDT < 50 as very low confidence, typically corresponding to disordered or poorly modeled regions (AlphaFold/EMBL-EBI guidelines). ColabFold also produced predicted aligned error (PAE) plots, which were inspected to evaluate the relative positioning of domains and to identify regions where inter-domain geometry is less reliable.

### PyMOL

In silico mutagenesis and three-dimensional protein structure analyses were performed using PyMOL (The PyMOL Molecular Graphics System, version 3.1.6.1; Schrödinger, LLC). The NLGN3 (Q9NZ94-2) sequence corresponding to the isoform expressed from the expression vector used in this study was modelled and analyzed as the “vector-derived” NLGN3 isoform, whereas the NLGN3 (AF-Q9NZ94-F1-v6) sequence was used as the AlphaMissense reference isoform because the exact vector-derived isoform was not available in the AlphaMissense resource. NLGN3 (Q9NZ94-2) and NLGN3 (AF-Q9NZ94-F1-v6) protein structures were superimposed using the “align” command in PyMOL, and backbone Cα root mean square deviation (RMSD) values were calculated to quantify structural differences between variants. In line with commonly used criteria in structural alignment and docking studies, RMSD values < 2.0 Å were interpreted as indicating highly similar conformations, values of 2.0–3.0 Å as moderate deviations, and values > 3.0 Å as substantial structural changes. During in silico mutagenesis, PyMOL’s mutagenesis wizard was also used to inspect local steric effects; large red discs were taken to indicate significant steric clashes between the introduced side chain and neighboring atoms, whereas small green or yellow indicators represented minor or near-contact interactions.

### AlphaMissense

Predictive pathogenicity scores for NLGN3 variants were obtained from the AlphaMissense dataset via the Hege Lab resource (https://alphamissense.hegelab.org/) ^21^. AlphaMissense pathogenicity scores range from 0 to 1, with higher scores indicating a greater likelihood of pathogenicity ^21,22^. In this study, we classified variants according to previously proposed thresholds as follows: variants with AlphaMissense scores ≥ 0.564 were considered likely pathogenic, those with scores between 0.34 and 0.564 as of uncertain significance, and those with scores < 0.34 as likely benign, following Cheng *et al.* ^31^ and subsequent benchmarking studies.

### Lolliplot

Lollipop plots illustrating the distribution of amino acid substitutions were generated in R (version 4.5.1) using the trackViewer package (version 1.44.0).

### TOGO var

For all variants analyzed, in silico pathogenicity prediction scores (SIFT, and PolyPhen**-**2) were retrieved from the Japanese genetic variation database TOGOVar (https://grch38.togovar.org/?mode=simple) ^32^. Both SIFT and PolyPhen**-**2 provide scores between 0 and 1 but with opposite directions of effect: lower SIFT scores indicate more deleterious variants, whereas higher PolyPhen**-**2 scores indicate more deleterious variants. In this study, SIFT scores ≤ 0.05 were considered deleterious, and PolyPhen–2 scores ≥ 0.85 were considered probably damaging.

### Plasmid constructs

All recombinant DNA experiments were performed in accordance with the guidelines established by the Institutional Biosafety Committee of Yamaguchi University and were approved under protocol number J22001.

All constructs were assembled in the pcDNA3.1(+) backbone under the control of the CMV promoter. The expression cassette comprises, in a single open reading frame from 5’ to 3’: the coding sequence of DsRed2, the I-CLiQ tag, and the coding sequence of human NLGN3 isoform 2 (UniProt: Q9NZ94-2), arranged as DsRed2-I-CLiQ-NLGN3. The native signal peptide of NLGN3 is retained at the N-terminus of the NLGN3 segment and therefore lies downstream of the DsRed2 and I-CLiQ sequences within the same open reading frame. The I-CLiQ tag is a modified form of the 2A peptide of Perina nuda virus. The parental 2A sequence is NIIGGGQKDLTQDGDVESNPGP ^33^; the sequence used here is NIIGGGQKDLTQHGDIESNPVF, which differs at four positions (D13H, V16I, G21V and P22F). The substitutions at positions 21 and 22 disrupt the NPGP motif required for ribosomal skipping, while the N-terminal region used as the antibody epitope is unchanged. The wild-type construct and all point mutants (NLGN3 WT, p.V341A ^25^, p.R451C ^34^, p.P514S ^25^, p.R597W ^35^, p.T632A ^36^) were custom synthesized and assembled by Uniqtech Inc. (Chiba, Japan). All constructs, including the element junctions and the introduced mutations, were verified by Sanger sequencing.

### Cell line

Human (female) embryonic kidney cells, HEK293T, were purchased from the RIKEN BioResource Center (RIKEN BRC; Ibaraki, Japan). HEK cells were cultured in Dulbecco’s Modified Eagle Medium (DMEM; 08459-64, Nacalai tesque, Inc., Kyoto, Japan) containing 10% fetal bovine serum (FBS; 35-079-C, Corning, NY, USA), 100 units/mL penicillin, and 100 μg/mL streptomycin (26253-84, Nakarai tesque Inc.); culture media was changed every two days. HEK cells were grown to subconfluence at 37°C in a humidified 5% CO_2_ incubator and passaged at a ratio of 1:4 - 1:8. Human neuroblastoma cells, SK-N-SH, were obtained from ATCC (HTB-11). SK-N-SH cells were cultured in Eagle’s Minimum Essential Medium (E-MEM; 051-07615, FUJIFILM Wako Pure Chemical Corporation, Osaka, Japan) containing 10% FBS (35-079-CV, Corning), 100 units/mL penicillin, and 100 μg/mL streptomycin; culture media was changed every two days. SK-N-SH cells were grown to subconfluence at 37°C in a humidified 5% CO_2_ incubator and passaged.

### I-CLiQ specific antibody

A custom rabbit polyclonal antibody specific for the I-CLiQ sequence was generated for this study (immunogen: HGDIESNP). The antibody was affinity-purified on a column of the immunizing peptide, and the purified stock was at 0.93 mg/mL. As part of the supplier’s quality control, the identity of the immunizing peptide was confirmed by mass spectrometry and the reactivity of the purified antibody against that peptide was confirmed by ELISA (enzyme-linked immunosorbent assay). In the cellular assay, non-transfected HEK293T cells stained under identical conditions showed no surface signal above background, indicating that the signal detected on transfected cells depends on expression of the I-CLiQ-tagged construct.

### Transient transfections

HEK cells were seeded at a density of 6 × 10^5^ cells/well in 6-well plates and incubated for 24 h. HEK cells were transfected with plasmid DNA (1.5 μg) using calcium chloride (120 μmol/mL, Sigma-Aldrich, Inc., St. Louis, MO, USA) in HEPES buffer (7365-45-9, Sigma-Aldrich, Inc.) with media change after 4 h of incubation, and collection 48 h later for flow cytometry.

For Polyethylenimine Max (PEI-MAX; 24765-100, Polysciences, Inc. PA, USA) -mediated transfection, HEK293T cells received 2 μg DNA at a DNA-to-PEI-MAX ratio of 1:2.5 (2 μg DNA: 5 μL PEI-MAX). SK-N-SH cells received 1 μg DNA at ratios of 1:3 and 1:5, or 2 μg DNA at ratios of 1:1.5, 1:2.5 and 1:3.5. For Lipofectamine 3000 (L3000001, Invitrogen, CA, USA)-mediated transfection, SK-N-SH cells received 4 μg DNA at a DNA-to-reagent ratio of 1:2.5 (4 μg DNA: 10 μL Lipofectamine 3000). All DNA-to-reagent ratios are expressed as μg DNA: μL reagent. In all cases, the medium was replaced 4 h after transfection, and cells were collected 48 h later for flow cytometry.

Transfection efficiency was evaluated separately using a DsRed-expressing vector, with the proportion of DsRed-positive cells quantified by a NovoCyte Flow Cytometer (Agilent, CA, USA).

### I-CLiQ Assay

Transfected HEK cells were trypsinized and placed on ice at 4°C for 10 min and then incubated with either anti-I-CLiQ antibody (rabbit polyclonal, 1:25 dilution of a 0.93 mg/mL stock, giving a final concentration of 37 µg/mL) in DMEM containing 2% FBS at 4°C for 2 h or an antibody against human NLGN3 (1:50; MAB6088, R&D Systems, Inc., MN, USA) in DMEM containing 2% FBS at 4°C for 1 h. Cells were washed with ice-cold PBS and incubated with Alexa Fluor 488 anti−Mouse IgG (1:250, A11011, Invitrogen,) or Alexa Fluor 488 anti-Rabbit IgG (1:250, A11034, Invitrogen) as secondary antibodies at 4°C for 30 min. Cells were subsequently washed with cold PBS and treated with membrane impermeant DAPI (340-07971, DOJINDO Laboratories, Kumamoto, Japan) to stain dead cells. Live cells were analyzed by flow cytometry using a NovoCyte Flow Cytometer, and the flow data was analyzed using NovoExpress software (Agilent). I-CLiQ-expression values for any given NLGN3^PMM^ are reported relative to NLGN3^WT^ and is a ratio of the calculated geometric mean fluorescence intensity (gMFI) of Alexa-488 that detects the extracellular NLGN3-I-CLiQ epitope relative to the gMFI of intracellular DsRed from the same group of live (DAPI negative) cells. I-CLiQ-assay values were calculated as follows: (gMFI Alexa-488 HEK-NLGN3^PMM^/gMFI DsRed HEK-NLGN3^PMM^)/ (gMFI Alexa-488 HEK-NLGN3^WT^/gMFI DsRed HEK-NLGN3^WT^). Values are reported as means ± SEM, as described under Quantification and statistical analysis.

For drug treatment analysis, transiently transfected HEK cells were plated at a density of 5 × 10^4^ cells/well into 6-well plates. After incubation for 24 h, cells were treated with either 4**-**phenylbutanoic acid (4**-**PBA; P21005, Sigma-Aldrich, Inc.) at a final concentration of 1 mM or 3-amino-1-propanesulfonic acid (3-APS; A76109, Sigma-Aldrich, Inc.) at a final concentration of 10 μM in DMEM with 2% FBS and incubated for a further 24 h. Cells were trypsinized for 1 min and antibody treatments were carried out prior to flow cytometry analysis.

### Western Blot

Cultured cells were washed with PBS and lysed with Mammalian Protein Extraction Reagent (M-PER, Thermo Fisher Scientific) with the ReadyShield Protease and Phosphatase Inhibitor Cocktail (PPC2020-1M, Sigma-Aldrich, Inc.). Protein samples were separated on a 4-15% SDS-PAGE gel (TGX gel, BioRad Laboratories, Inc., CA, USA) and transferred to polyvinylidene fluoride (PVDF) membrane. Membranes were incubated in Block Ace (UK-B80; BioRad Laboratories, Inc.) at 4°C overnight and then with an antibody against human NLGN3 (1:1,000; MAB6088, R&D Systems, Inc.) and human GAPDH (1:5,000; CB1001, Merck Millipore, MA, USA) as primary monoclonal antibodies. Membranes were washed three times in PBS with 0.1% Tween-20, then with horseradish peroxidase (HRP)-conjugated anti-mouse IgG (1:5,000; NA931V, GE Healthcare, IL, USA) as secondary antibodies. All antibody incubations were performed at room temperature (RT) for 1-hr. Blots were developed using ImmunStar LD Chemiluminescence Kit (292-69903, FUJIFILM Wako Pure Chemical Corporation) and imaged using an Amersham Imager (GE Healthcare).

### Immunofluorescence

Transfected HEK293T cells were fixed in 4% Paraformaldehyde (PFA; Nacalai tesque, Inc.) for 10 min at 4°C and washed in PBS. No permeabilization step was included, so that only surface-exposed NLGN3 was accessible to the antibody. Cells were blocked with 3% bovine serum albumin (BSA, Sigma-Aldrich, Inc.) at RT for 1 h and then incubated with a primary antibody against human NLGN3 (1:200) at RT for 1 h. After wash with PBS, cells were incubated with Alexa Fluor 488 anti-mouse IgG (Thermo Fisher Scientific) as a secondary antibody at RT for 1 h. Coverslips were enclosed using ProLong Glass Antifade Mount (Thermo Fisher Scientific) after DAPI treatment at RT for 5 min. Images were obtained with a BZ-9000 or BZ-X1000 (Keyence, Osaka, Japan).

### Quantification and statistical analysis

Statistical analyses were performed in R (version 4.5.1). Data are presented as means +/- standard error of the mean (SEM), and the number of independent experiments is given in each figure legend. For comparisons among three or more groups, one-way analysis of variance (ANOVA) was used, followed by Tukey’s honestly significant difference (HSD) post hoc test for pairwise comparisons. For comparisons between two groups, Student’s unpaired two-tailed t-test was used. Two-sided *p* values below 0.05 were considered statistically significant.

## Acknowledgement

We thank Ms. Saki Horie for their excellent technical assistance with the experiments. We are also grateful to the members of the first term of the ACT-X Project (JST) and our mentor for their valuable discussions. We thank the Science Research Center of Yamaguchi University, including the Institute of Gene Research, the Institute for Biomedical Research, and the Institute of Life Science and Medicine, for their support. This work was conducted using research equipment shared through the MEXT Project for Promoting Public Utilization of Advanced Research Infrastructure (Program for Supporting Construction of Core Facilities), Grant Number JPMXS0440400024.

## Author contribution

N.T. conceived and designed the study. N.T. and K.T. developed the I-CLiQ methodology. N.T. performed the investigation, analyzed the data, and created the visualizations. R.O. provided technical assistance with the experiments and helped with manuscript preparation. N.T. wrote the original draft of the manuscript. N.T., K.T., and T.Y. contributed to the writing, review, and editing of the final manuscript. All authors have read and approved the published version of the manuscript.

## Data and materials availability

The datasets supporting the findings of this study are available from the following public repositories: AlphaFold Protein Structure Database (https://alphafold.ebi.ac.uk/), AlphaMissense via Hege Lab (https://alphamissense.hegelab.org/), ColabFold (https://colab.research.google.com/github/sokrypton/ColabFold/blob/main/AlphaFold2.ipynb#scrollTo=ADDuaolKmjGW), TogoVar (https://grch38.togovar.org/?mode=simple), NCBI (https://www.ncbi.nlm.nih.gov/gene/54413), and Ensembl (http://www.ensembl.org/Homo_sapiens/Gene/Summary?db=core;g=ENSG00000196338;r=X:71143286-71175255). Primary data underlying the figures, including raw flow cytometry data files, uncropped immunoblot images, original immunofluorescence images, and the R scripts used for analysis, are available from N.T. upon reasonable request. Requests should be submitted to.

