## Supplementary Information for "NLGN3-tagging system for Evaluating Membrane Localization Applied to Drug Screening"

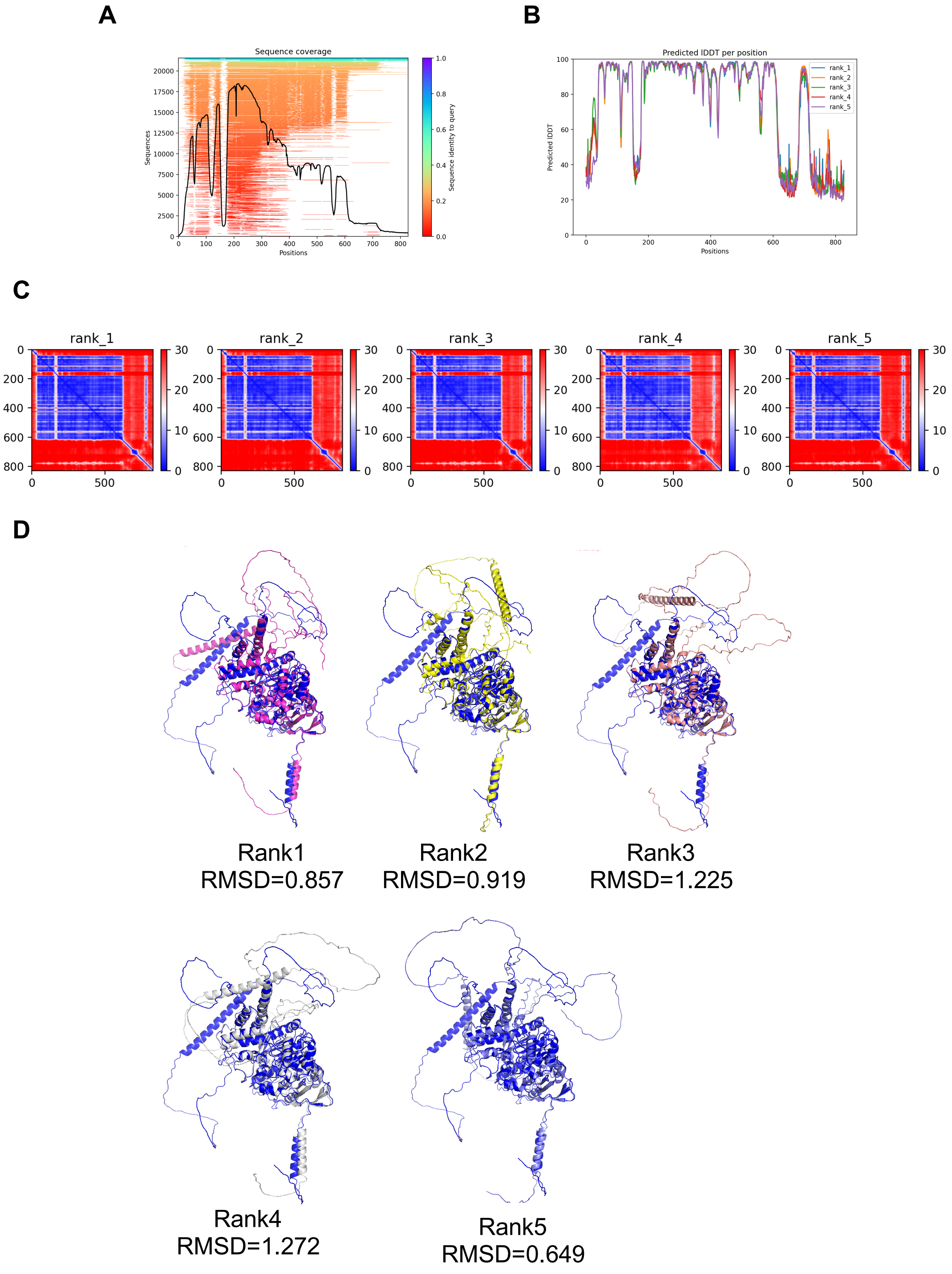

A
Sequence coverage
Sequences
Positions
Sequence identity to query
B
Predicted lDDT per position
Predicted lDDT
Positions
rank_1
rank_2
rank_3
rank_4
rank_5
C
rank_1
rank_2
rank_3
rank_4
rank_5
D
Rank1
RMSD=0.857
Rank2
RMSD=0.919
Rank3
RMSD=1.225
Rank4
RMSD=1.272
Rank5
RMSD=0.649

**Figure S1. Structural analysis of two NLGN3 isoforms.**

Pairwise structural alignment of NLGN3 (Q9NZ94-2, experimental target) and NLGN3 (AF-Q9NZ94-F1-v6, a model from AlphaFold). **(A)** Sequence coverage of the MSA for Q9NZ94-2. **(B)** Predicted lDDT values along the sequence of Q9NZ94-2, as predicted by ColabFold. **(C)** Predicted Aligned Error (PAE) plot for the top five ColabFold-predicted NLGN3 models (ranks 1–5), demonstrating high confidence in domain positioning. **(D)** Superimposition of the ColabFold-predicted structures of Q9NZ94-2 (ranks 1–5, colored) and AF-Q9NZ94-F1-v6 (gray). The RMSD values between the two structures were 0.857 Å (rank 1), 0.919 Å (rank 2), 1.225 Å (rank 3), 1.272 Å (rank 4), and 0.649 Å (rank 5).

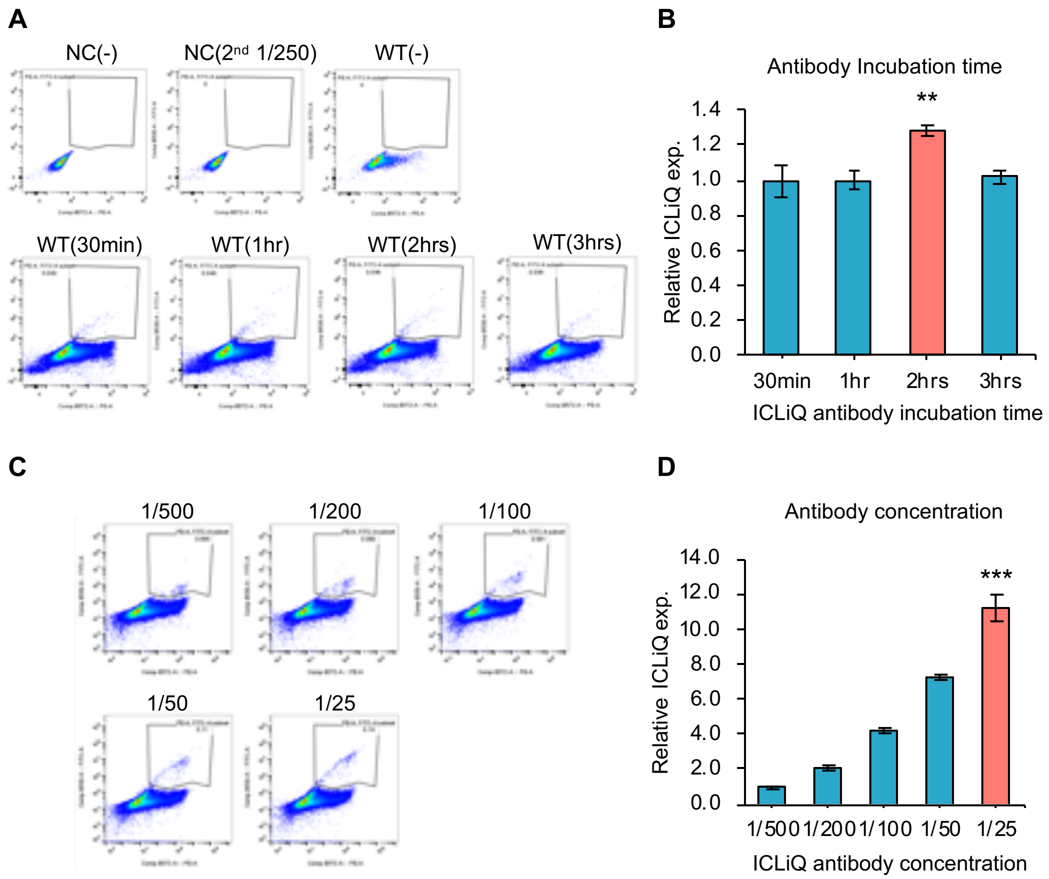

**Figure S2. Optimization of antibody incubation time and concentration for live-cell surface staining.**

**(A)** the representative density plots of panel B, and **(B)** Bar graph shows the mean fluorescence intensity. The signal intensity reached a plateau at 2 hours. All pairwise comparisons between "2 hrs" and each of the other groups yielded $**P$ values $< 0.01$. **(C)** the representative density plots of panel D, and **(D)** Bar graph shows the mean fluorescence intensity. All pairwise comparisons between "1/25" and each of the other groups yielded. $***$ $P$ values $< 0.001$. Data are presented as mean ± S.E.M.

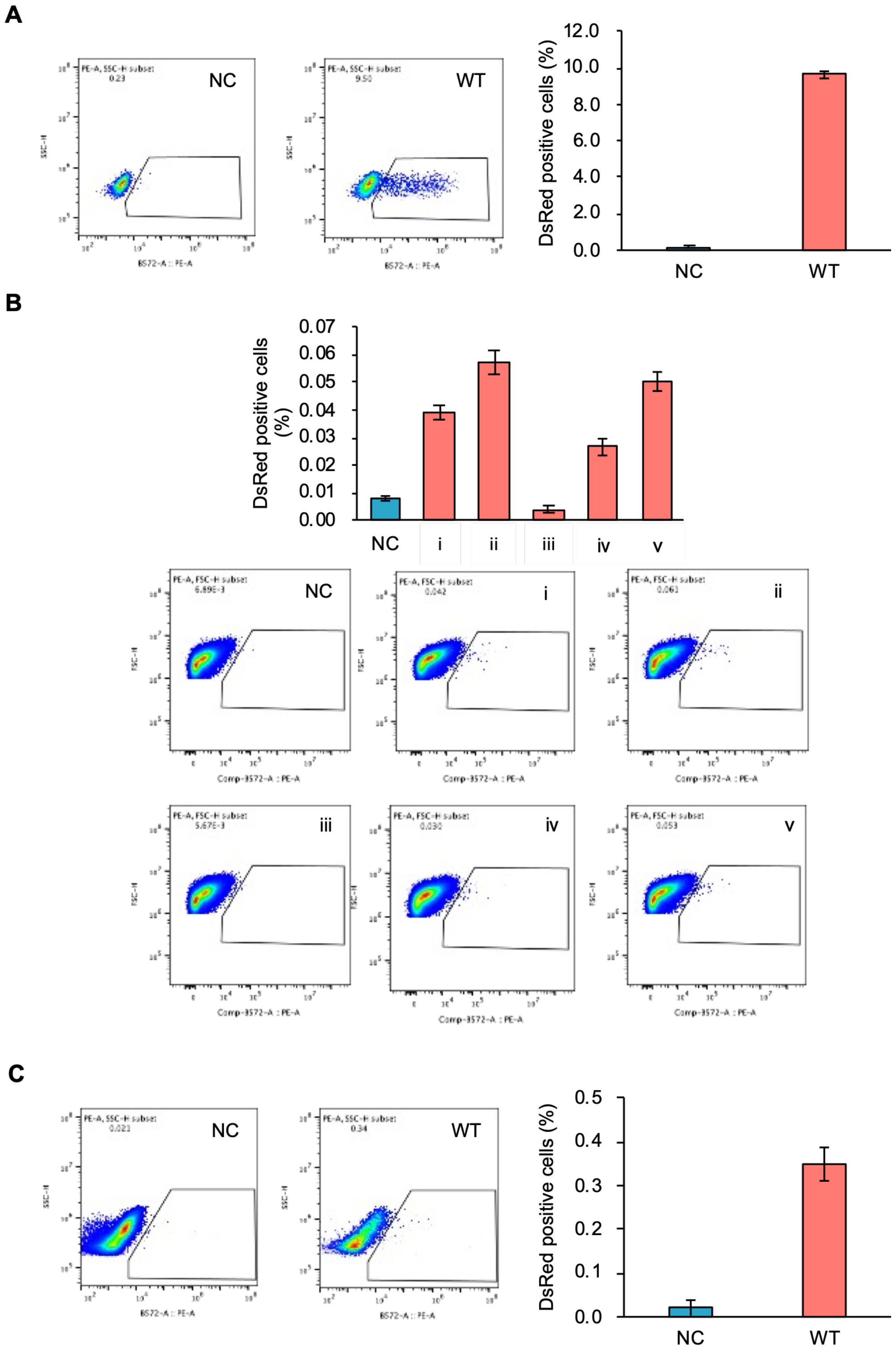
A
NC
WT
DsRed positive cells (%)
12.0
10.0
8.0
6.0
4.0
2.0
0.0
NC
WT
B
DsRed positive cells (%)
0.07
0.06
0.05
0.04
0.03
0.02
0.01
0.00
NC
i
ii
iii
iv
v
NC
i
ii
iii
iv
v
C
NC
WT
DsRed positive cells (%)
0.5
0.4
0.3
0.2
0.1
0.0
NC
WT

**Figure S3. Transfection of ICLiQ-WT vector into HEK293T and SK-N-SH cells.**

Transfection efficiency was evaluated by flow cytometric quantification of DsRed-positive cells following introduction of a DsRed-expressing vector. All DNA-to-reagent ratios are expressed as μg DNA : μL reagent. NC, no-DNA control. **(A)** HEK293T cells transfected with the DsRed-expressing vector using PEI-MAX at a DNA:PEI-MAX ratio of 1:2.5 (2 μg DNA : 5 μL PEI-MAX), analyzed by flow cytometry. **(B)** SK-N-SH cells transfected with the DsRed-expressing vector using PEI-MAX at the indicated DNA:PEI-MAX ratios: (i) 1:3, (ii) 1:5, (iii) 1:1.5, (iv) 1:2.5, and (v) 1:3.5. Conditions (i) and (ii) were performed with 1 μg DNA; conditions (iii)–(v) were performed with 2 μg DNA. **(C)** SK-N-SH cells transfected with the DsRed-expressing vector using Lipofectamine 3000 at a DNA:Lipofectamine 3000 ratio of 1:2.5 (4 μg DNA : 10 μL Lipofectamine 3000).

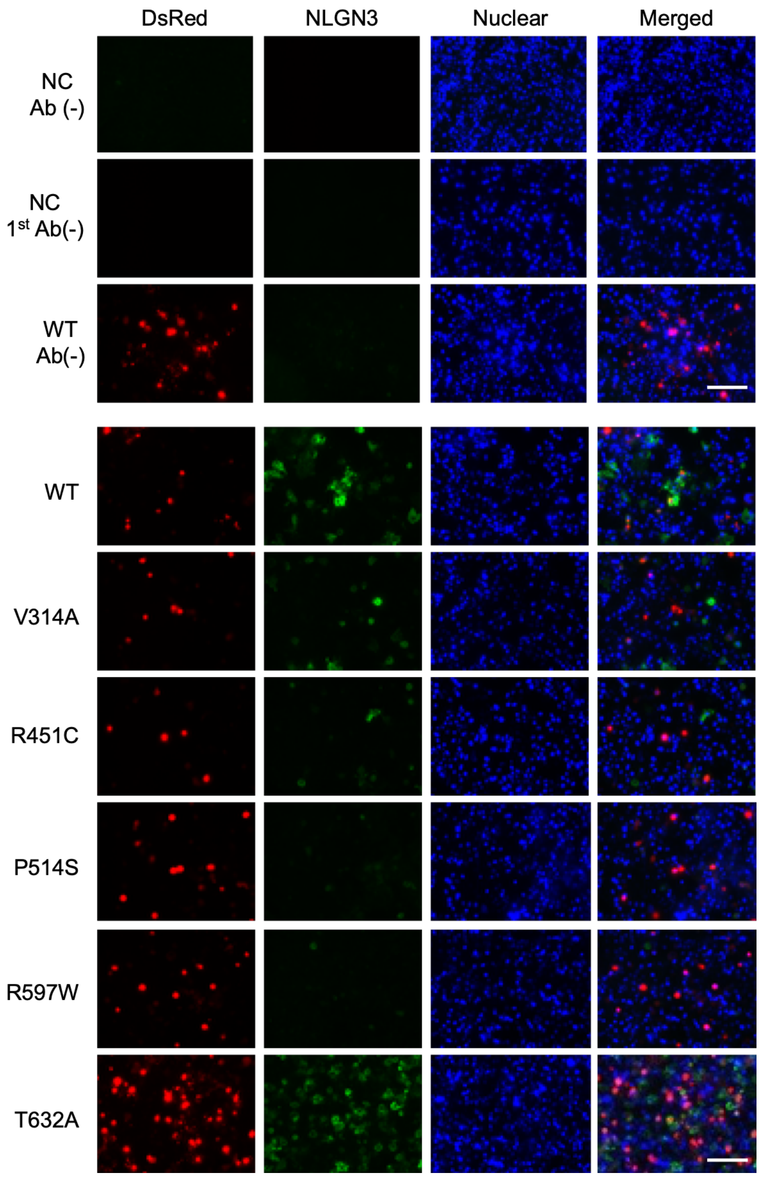

**Figure S4. Immunofluorescence staining of NLGN3.**

HEK293T cells were processed under identical conditions, except that the primary antibody or the expression vector was omitted. (NC, Ab(−)) Non-transfected cells without antibody. (NC, 1st Ab(−)) Cells incubated with the secondary antibody only (no primary antibody). (WT, Ab(−)) WT vector–transfected cells without antibody. No specific fluorescence signal was detected under any of these conditions, confirming that the observed signals are specific to NLGN3 expression. Low-magnification (40× objective) images are shown. Scale bar shows 100 μm.

**Table S1. NLGN3 isoform alignment.**

| | | | |
|---|---|---|---|
| Query: NLGN3 AF-Q9NZ94-F1-v6 sequence | | | |
| Subject: NLGN3 Q9NZ94-2 sequence | | | |
| Gaps:20/848(2%) | | | |
| Query | 1 | MWLRLGPPSLSLSPKPTVGRSLCLTLWFLSLALRASTQAPAPTVNTHFGKLRGARVPLPS | 60 |
| | | MWLRLGPPSLSLSPKPTVGRSLCLTLWFLSLALRASTQAPAPTVNTHFGKLRGARVPLPS | |
| Subject | 1 | MWLRLGPPSLSLSPKPTVGRSLCLTLWFLSLALRASTQAPAPTVNTHFGKLRGARVPLPS | 60 |
| Query | 61 | EILGPVDQYLGVPYAAPPIGEKRFLPPEPPPSWSGIRNATHFPPVCPQNIHTAVPEVMLP | 120 |
| | | EILGPVDQYLGVPYAAPPIGEKRFLPPEPPPSWSGIRNATHFPPVCPQNIHTAVPEVMLP | |
| Subject | 61 | EILGPVDQYLGVPYAAPPIGEKRFLPPEPPPSWSGIRNATHFPPVCPQNIHTAVPEVMLP | 120 |
| Query | 121 | VWFTANLDIVATYIQEPNEDCLYLNVYVPTEDVKRISKECARKPNKKICRKGGSGAKKQG | 180 |
| | | VWFTANLDIVATYIQEPNEDCLYLNVYVPTED GSGAKKQG | |
| Subject | 121 | VWFTANLDIVATYIQEPNEDCLYLNVYVPTED--------------------------------------GSGAKKQG | 160 |
| Query | 181 | EDLADNDGDEDEDIRDSGAKPVMVYIHGGSYMEGTGNMIDGSILASYGNVIVITLNYRVG | 240 |
| | | EDLADNDGDEDEDIRDSGAKPVMVYIHGGSYMEGTGNMIDGSILASYGNVIVITLNYRVG | |
| Subject | 161 | EDLADNDGDEDEDIRDSGAKPVMVYIHGGSYMEGTGNMIDGSILASYGNVIVITLNYRVG | 220 |
| Query | 241 | VLGFLSTGDQAAKGNYGLLDQIQALRWVSENIAFFGGDPRRITVFGSGIGASCVSLLTLS | 300 |
| | | VLGFLSTGDQAAKGNYGLLDQIQALRWVSENIAFFGGDPRRITVFGSGIGASCVSLLTLS | |
| Subject | 221 | VLGFLSTGDQAAKGNYGLLDQIQALRWVSENIAFFGGDPRRITVFGSGIGASCVSLLTLS | 280 |
| Query | 301 | HHSEGLFQRAIIQSGSALSSWAVNYQPVKYTSLLADKVGCNVLDTVDMVDCLRQKSAKEL | 360 |
| | | HHSEGLFQRAIIQSGSALSSWAVNYQPVKYTSLLADKVGCNVLDTVDMVDCLRQKSAKEL | |
| Subject | 281 | HHSEGLFQRAIIQSGSALSSWAVNYQPVKYTSLLADKVGCNVLDTVDMVDCLRQKSAKEL | 340 |
| Query | 361 | VEQDIQPARYHVAFGPVIDGDVIPDDPEILMEQGEFLNYDIMLGVNQGEGLKFVEGVVDP | 420 |
| | | VEQDIQPARYHVAFGPVIDGDVIPDDPEILMEQGEFLNYDIMLGVNQGEGLKFVEGVVDP | |
| Subject | 341 | VEQDIQPARYHVAFGPVIDGDVIPDDPEILMEQGEFLNYDIMLGVNQGEGLKFVEGVVDP | 400 |
| Query | 421 | EDGVSGTDFDYSVSNFVDNLYGYPEGKDTLRETIKFMYTDWADRDNPETRRKTLVALFTD | 480 |
| | | EDGVSGTDFDYSVSNFVDNLYGYPEGKDTLRETIKFMYTDWADRDNPETRRKTLVALFTD | |
| Subject | 401 | EDGVSGTDFDYSVSNFVDNLYGYPEGKDTLRETIKFMYTDWADRDNPETRRKTLVALFTD | 460 |
| Query | 481 | HQWVEPSVVTADLHARYGSPTYFYAFYHHCQSLMKPAWSDAAHGDEVPYVFGVPMVGPTD | 540 |
| | | HQWVEPSVVTADLHARYGSPTYFYAFYHHCQSLMKPAWSDAAHGDEVPYVFGVPMVGPTD | |
| Subject | 461 | HQWVEPSVVTADLHARYGSPTYFYAFYHHCQSLMKPAWSDAAHGDEVPYVFGVPMVGPTD | 520 |
| Query | 541 | LFPCNFSKNDVMLSAVVMTYWTNFAKTGDPNKPVPQDTKFIHTKANRFEEVAWSKYNPRD | 600 |
| | | LFPCNFSKNDVMLSAVVMTYWTNFAKTGDPNKPVPQDTKFIHTKANRFEEVAWSKYNPRD | |
| Subject | 521 | LFPCNFSKNDVMLSAVVMTYWTNFAKTGDPNKPVPQDTKFIHTKANRFEEVAWSKYNPRD | 580 |
| Query | 601 | QLYLHIGLKPRVRDHYRATKVAFWKHLVPHLYNLHDMFHYTSTTTKVPPPDTTHSSHITR | 660 |
| | | QLYLHIGLKPRVRDHYRATKVAFWKHLVPHLYNLHDMFHYTSTTTKVPPPDTTHSSHITR | |
| Subject | 581 | QLYLHIGLKPRVRDHYRATKVAFWKHLVPHLYNLHDMFHYTSTTTKVPPPDTTHSSHITR | 640 |
| Query | 661 | RPNGKTWSTKRPAISPAYSNENAQGSWNGDQDAGPLLVENPRDYSTELSVTIAVGASLLF | 720 |
| | | RPNGKTWSTKRPAISPAYSNENAQGSWNGDQDAGPLLVENPRDYSTELSVTIAVGASLLF | |
| Subject | 641 | RPNGKTWSTKRPAISPAYSNENAQGSWNGDQDAGPLLVENPRDYSTELSVTIAVGASLLF | 700 |
| Query | 721 | LNVLAFAALYYRKDKRRQEPLRQPSPQRGAGAPELGAAPEEELAALQLGPTHHECEAGPP | 780 |
| | | LNVLAFAALYYRKDKRRQEPLRQPSPQRGAGAPELGAAPEEELAALQLGPTHHECEAGPP | |
| Subject | 701 | LNVLAFAALYYRKDKRRQEPLRQPSPQRGAGAPELGAAPEEELAALQLGPTHHECEAGPP | 760 |
| Query | 781 | HDTLRLTALPDYTLTLRRSPDDIPLMTPNTITMIPNSLVGLQTLHPYNTFAAGFNSTGLP | 840 |

| | | | |
|---|---|---|---|
| | | HDTLRLTALPDYTLTLRRSPDDIPLMTPNTITMIPNSLVGLQTLHPYNTFAAGFNSTGLP | |
| Subject | 761 | HDTLRLTALPDYTLTLRRSPDDIPLMTPNTITMIPNSLVGLQTLHPYNTFAAGFNSTGLP | 820 |
| Query | 841 | HSHSTTRV | 848 |
| | | HSHSTTRV | |
| Subject | 821 | HSHSTTRV | 828 |
